# CellTrap: A Microfluidic Platform Enabling Cell-Cell Interactions at Variable Effector to Target Ratios

**DOI:** 10.64898/2025.12.25.696500

**Authors:** Muhammad Zia Ullah Khan, Morteza Hasanzadeh Kafshgari, Ali Bashiri Dezfouli, Oliver Hayden, Gabriele Multhoff, Ghulam Destgeer

## Abstract

Immune–cancer cell interactions play a central role in understanding antitumor responses and evaluating immunotherapies. However, long-term, single-cell–level analysis of these interactions remains challenging. To address this, we developed a microfluidic trapping device with 1,024 traps, each equipped with a filter to retain cells, sustain medium flow, minimize cross-talk, and allow precise control of effector-to-target (E:T) ratios. The platform enables continuous monitoring of immune–cancer interactions for up to 14 hours. Device characterization was performed using 10 µm fluorescent beads seeded via hydrostatic flow, with trap occupancy validated by Poisson statistics. Initial experiments using PBMCs against GFP-expressing U87 (U87^GFP^) glioblastoma cells demonstrated an immune-mediated reduction in GFP intensity, which was interpreted cautiously as a cytotoxic response. To improve reproducibility, we subsequently employed IL-2-stimulated Natural Killer cells (NK92^IL2^) as standardized effectors and evaluated their interactions with U87^GFP^ glioblastoma cells, K562 chronic myelogenous leukemia cells, and LS174T adenocarcinoma cells. Time-lapse imaging revealed transient intracellular calcium fluxes, consistent with early activation of NK92^IL2^ cells, followed by a cytotoxic response. Increasing E:T ratios consistently enhanced immune activity, highlighting the utility of this device for dissecting immune–cancer interactions and guiding the development of immunotherapy.

## 1. Introduction

Cell-to-cell interactions are crucial for understanding the complexity of immune responses. They underpin our understanding of tumor survival mechanisms^1^, drive the development of advanced immunotherapies such as CAR T-cell and CAR NK-cell therapies, enable personalized treatment strategies^2^, and inform efforts to prevent metastasis^3^.

To investigate these intricate immune dynamics, researchers employ a range of *in vivo*, *ex vivo*, and *in vitro* methodologies, each offering distinct strengths and facing specific limitations^4^. *In vivo* models provide the most biologically comprehensive insights but are often constrained by ethical considerations, technological challenges, extended timelines, and high costs. *Ex vivo* approaches, which rely on biopsies or intact tissues, enable the direct examination of human or animal samples; however, they are limited by the restricted availability of patient material and issues such as tissue degradation over time.

In contrast, *in vitro* systems offer accessibility and scalability, making them a mainstay in immunological research. Yet, conventional in vitro platforms, such as Petri dishes and well plates, lack fine control over the cellular microenvironment. As a result, they tend to average out cellular responses, obscuring critical single-cell interactions within heterogeneous populations. Moreover, their dependence on large cell quantities hinders the study of rare primary cells or genetically engineered immune subsets.

Microfluidics-based platforms have emerged as a promising alternative, addressing many of these limitations by enabling high-resolution analysis^5^ and real-time observation of cell behavior and interactions^6,7^. These systems offer precise control over the microenvironment, i.e., regulating chemical gradients, cell numbers, and interaction timeframes, while requiring only minimal sample and reagent volumes^8,9^. Such features make microfluidic technologies highly attractive for applications in immunotherapy^10^ and personalized medicine^11^.

A variety of microfluidic architectures have been developed to study cell–cell interactions, including microwells^12–22^, droplet encapsulation systems^23–29^, and array-based trapping platforms^30,31^. These microfluidic cell-pairing strategies can be broadly categorized into deterministic and stochastic approaches^32^. Deterministic methods, which can be further categorized into passive and active mechanisms, enable controlled cell pairing and achieve high one-to-one pairing efficiencies. Passive deterministic systems typically rely on size-based cell trapping and demand precisely engineered microstructures to ensure proper alignment and capture. Active deterministic approaches, on the other hand, employ external forces such as electric, magnetic, acoustic, or optical fields to manipulate cells. While highly effective, these systems often require expensive instrumentation, specialized expertise, and substantial physical space. These factors hinder their widespread use in routine biological research. In contrast, stochastic pairing, governed by Poisson statistics, offers a simple and scalable alternative. It accommodates a broad range of cell sizes and simplifies device design, albeit at the cost of lower one-to-one pairing efficiency. Despite these trade-offs, both deterministic and stochastic platforms have provided critical insights into immune–cancer cell interactions, enabling analyses such as time-lapse imaging, calcium flux monitoring, cytokine release assays, and cytotoxicity measurements across a diverse range of E:T ratios.

Nonetheless, each of these microfluidic architectures entails specific trade-offs. Microwell and microarray-based systems facilitate medium exchange, staining, and cell adhesion; however, they are prone to crosstalk between adjacent wells or traps and typically require high cell concentrations to achieve sufficient occupancy. Droplet microfluidic systems, by contrast, effectively prevent crosstalk but preclude medium exchange and cell adhesion. They also involve technically complex setups, relying on oil emulsions, stabilization devices for imaging, and pre-mixing of staining reagents, which can introduce cytotoxic effects. Moreover, both microwell and droplet-based systems often rely on expensive syringe pumps, which further increases system complexity and cost. Together, these limitations highlight the need for a microfluidic platform that leverages the strengths of existing systems while mitigating their drawbacks. Specifically, this requires a device that enables rapid and controlled cell seeding, supports medium exchange without crosstalk, operates instrument-free, and accommodates small sample volumes for rare or engineered cell populations.

To meet these requirements, we have developed a polydimethylsiloxane (PDMS)-based microfluidic platform, featuring 1,024 parallel-arranged traps in a 1D array, specifically designed for long-term co-incubation and real-time visualization of hundreds of cell–cell interactions (**Figure 1**). The gas-permeable PDMS channel enables sustained culture and live-cell imaging using standard microscopes equipped with environmental control. The platform supports instrument-free sample loading through hydrostatic pressure-driven flow, ensuring robust operation and enabling efficient medium exchange with minimal crosstalk (Figure 1A).

**Figure 1.**
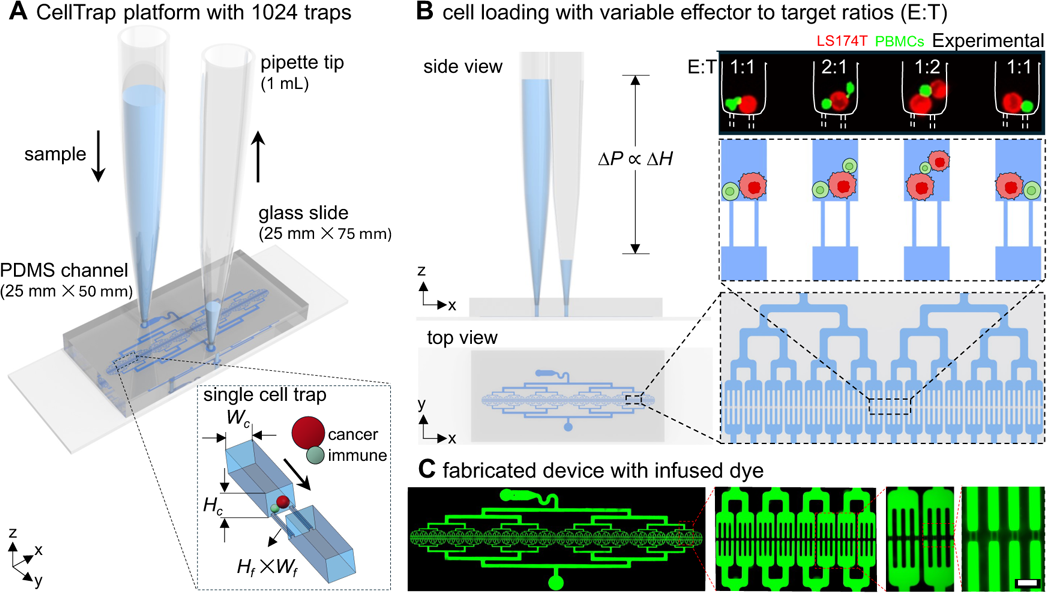
An instrument-free operation of the CellTrap platform. (A) A PDMS microfluidic channel containing 1,024 traps is bonded to a glass slide. Sample loading is driven by hydrostatic pressure generated by the fluid column at the inlet pipette tip. The main channel height (*H_c_*) and width (*W_c_*) are larger than the cell diameter (*D_c_*) to allow uninterrupted cell flow, whereas the filter height (*H_f_*) and width (*W_f_*) at each trap are smaller than *D_c_* to ensure cell capture. (B) The hydrostatic pressure gradient (Δ*P*) is proportional to the height difference between the inlet and outlet free surfaces. Cells are trapped at variable E:T ratios, e.g., 1:1, 2:1, 1:2, and 1:1. (C) Fabricated device infused with Rhodamine B dye to highlight the channel network. Scale bar: 50µm.

The channel geometry is optimized for selective trapping: the main channel height (*H_c_*) and width (*W_c_*) exceed the average cell diameter (*D_c_*) to facilitate smooth flow, while the filter height (*H_f_*) and width (*W_f_*) at the trap are smaller than *D_c_* to capture cells effectively. For example, a single cancer cell (red) paired with an engineered immune cell (green) can be co-incubated for several days, allowing quantitative time-lapse analysis of immune efficacy. Each trap is capable of stochastically capturing two distinct cells, generating controlled pairings across a natural spectrum of E:T ratios (Figure 1B).

The CellTrap platform is fabricated with eight bifurcation stages highlighted by the infusion of channels with a fluorescent dye (Figure 1C). At the final stage of bifurcation, the main channel splits into four parallel daughter channels that terminate at the cell trap, each with two to three filter channels to capture cancer and immune cells. We have characterized our CellTrap platform for stochastic loading of cells. As a proof of concept, we first performed cytotoxicity assays to investigate interactions between U87^GFP^ cells and peripheral blood mononuclear cells (PBMCs). Secondly, we performed calcium and cytotoxicity assays to investigate the interactions of three cancer cell lines, i.e., glioblastoma U87^GFP^, chronic myelogenous leukemia K562, and adenocarcinoma LS174T, against NK92^IL2^ immune cells at variable E:T ratios. We have selected these cancer cell types for their diverse range and levels of ligand expression that activate or inhibit the response of NK cells.

## 2. Results and discussion

### 2.1 Stochastic loading of traps to achieve variable E:T ratios

The trapping of particles or cells in the CellTrap platform follows a Poisson distribution due to the stochastic nature of their flow. First, we characterized the CellTrap device using 10-µm fluorescent microparticles and subsequently validated the device with cells (**Figure 2**). We evaluated the device’s performance numerically and experimentally under various operating conditions, including pump versus pipette loading, mixed versus non-mixed samples, and different filter numbers (**Figure S1**). No significant deviation from the analytical Poisson distribution was observed under any condition (**Figure S2**). Therefore, pipette-based loading with pre-mixed samples was selected for subsequent experiments due to its operational simplicity and reduced risk of cell damage.

**Figure 2.**
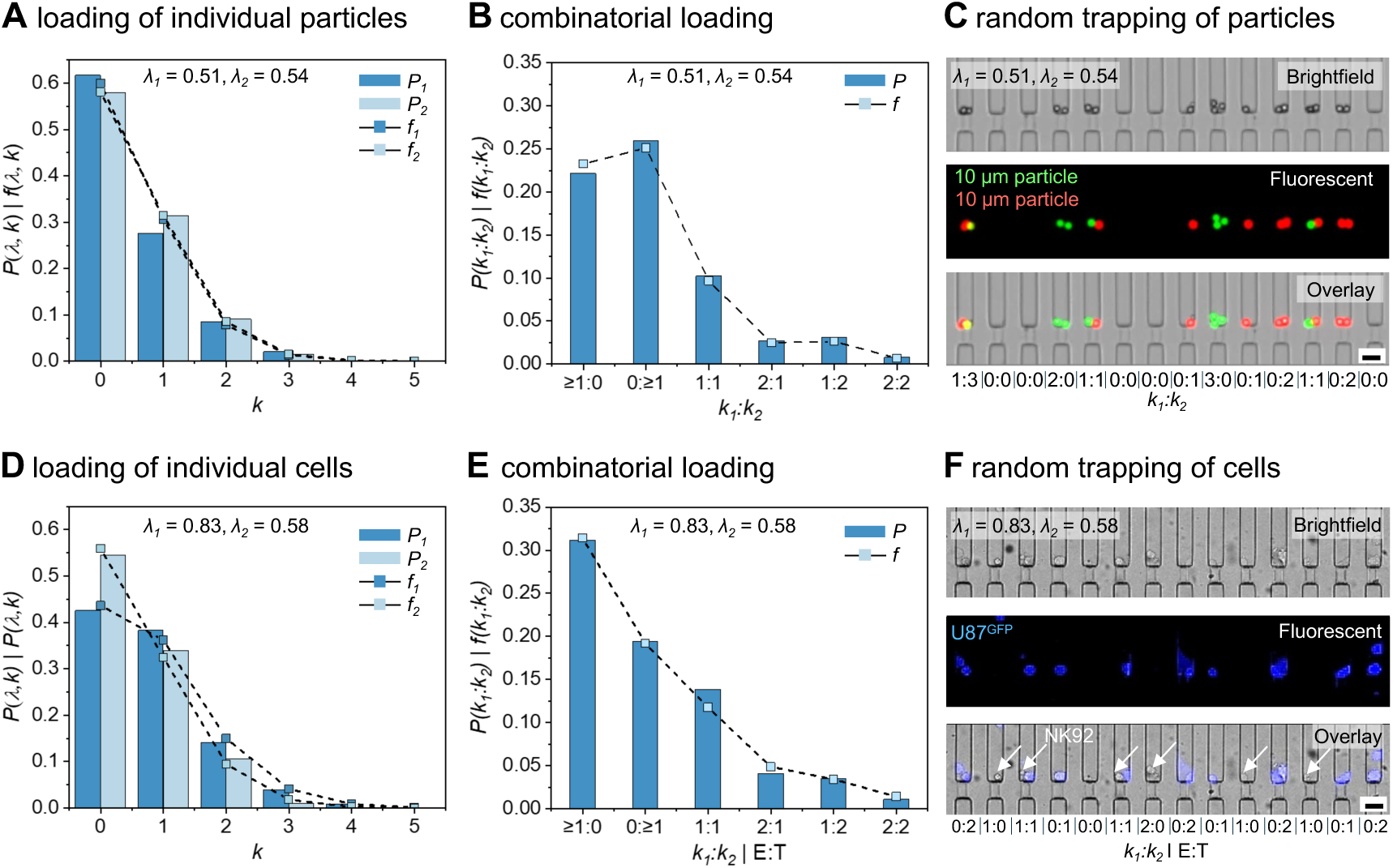
Characterization of the CellTrap Device with 1,024 Traps. (A) The experimental loading frequencies (*f*) of individual particles, i.e., green and red, are compared to the theoretical Poisson distributions (*P*) for *λ_1_* = 0.51 (green) and *λ_2_* = 0.54 (red). (B) A combinatorial loading distribution of the two particle types was analyzed using a double Poisson distribution, which is compared with the experimental dual loading frequency. (C) Representative bright-field, fluorescence, and overlay images of red and green particles trapped in a channel at different *k_1_*:*k_2_* ratios. (D) The experimental loading frequencies (*f*) and the theoretical Poisson distributions (*P*) are plotted for the U87^GFP^ and NK92^IL2^ cells seeded in the CellTrap device with *λ_1_* = 0.83 (U87^GFP^) and *λ_2_* = 0.58 (NK92^IL2^). (E) The experimental dual loading frequency and theoretical double Poisson distributions result in varying E:T or *k_1_*:*k_2_* ratios. (F) Representative bright-field, fluorescence, and overlay images of cancer (purple) and immune (white arrows) cells trapped inside a channel at different E:T ratios. (scale bars: 30 μm)

Sample loading can be adjusted by varying the initial sample concentration and seeding duration, which determine the distribution of trapped particles or cells across the 1,024 traps. After the sample loading, an average number of particles per trap (*λ = N_p_*/*N_t_*) is measured for each experiment, where *N_p_* is the total number of trapped particles in the device and *N_t_* (= 1,024) is the total number of traps. This *λ* value is used to evaluate the analytical Poisson distribution *P*(*λ, k*) to compare with the experimental frequency *f* (*λ, k*) of loading events for a given trap (*k*). The theoretical probability of *k* for a known *λ* is given by *P*(*λ, k*) = (*λ^k^ e^-λ^ / k!*), where *k* = 0, 1, 2, … is the number of particles in one trap. The experimental frequency of *k* is calculated as *f*(*λ, k*) = *n_t_*(*k*)/*N_t_*, where *n_t_* is the number of traps for a given *k =* 0, 1, 2, ….

We seeded a mixture of two particles, i.e., green (1) and red (2), into the CellTrap device and measured their experimental loading frequencies, *f_1_* and *f_2_*, respectively, across the 1,024 traps, which aligned well with the theoretical probabilities (*P_1_* and *P_2_*) for both particles, respectively (Figure 2A-2C). Here, a *λ_1_* = 0.51 resulted in more than half of the traps empty of green particles (i.e., *n_t,1_*(0) = 632, *f* (0) = 62%), whereas 28% of the traps had a single green particle loaded (i.e, *n_t,1_*(1) = 283), matching the theoretical predictions (Figure 2A). The remaining 10% of the traps were occupied by two or more green particles per trap. A similar trend is observed for the red particles with *λ_2_* = 0.54. By varying *λ*, the theoretical probabilities for different values of *k* can be evaluated (Figure S3). When the experimentally observed frequencies of *k* agree with these theoretical probabilities, the experimental distribution can be predicted and tuned both for single and multiple loadings, according to the theoretical trend.

A combinatorial loading distribution of the two particle types was analyzed using a double Poisson distribution, defined as: *P*(*k_1_*:*k_2_*) = *P*(*k_1_*) × *P*(*k_2_*) for corresponding means *λ_1_* and *λ_2_* (Figure 2B). For comparison, the experimental dual loading frequency was determined as: *f*(*k_1_*:*k_2_*) = *f*(*k_1_*) × *f*(*k_2_*). Approximately 10% of the traps contained exactly one particle of each type, i.e., *f*(1:1) ≈ 0.1, corresponding to *n_t_*(1:1) = 105. Traps with *k_1_*:*k_2_* = 1:2 and 2:1 each accounted for approximately 3% of the total traps, i.e., *n_t_*(1:2) = 32, *n_t_*(2:1) = 30. A small fraction of traps (< 2%) contained multiplets with *k_1_*:*k_2_* = (≥2:≥2). A majority of the traps ≈ 38% were empty ( i.e., *k_1_*:*k_2_* = 0:0, *n_t_*(0:0) = 391). The remaining traps with *k_1_*:*k_2_* = 0:≥1 (*n_t_* = 241) and ≥1:0 (*n_t_* = 202) were accounted as control groups. The experimental value of *f*(1:1) = 10% matched the theoretical prediction of 9.2% double (1:1) occupancies using *λ_1,2_* = 0.5 by the Poisson distribution (**Figure S3**). Experimental images of the CellTrap devices confirmed these variable *k_1_*:*k_2_* ratios of the loaded particles (Figure 2C). Diverse *k_1_*:*k_2_* ratios were achieved within the same device, enabling a mixed population of empty (0:0), control (0:≥1 or ≥1:0), multiplet (≥1:≥1, except 1:1), and singleton (1:1) traps, with their proportions adjustable by varying *λ*.

We next extended this analysis to living cells. U87^GFP^ (*λ_1_* = 0.83) and NK92^IL2^ (*λ_2_* = 0.58) cells exhibited similar stochastic loading behavior (Figure 2D). Approximately 13.8% of the traps contained exactly one cell of each type, i.e., *f*(1:1) ≈ 0.138, corresponding to *n_t_*(1:1) = 142, closely matching the theoretical prediction of 11.8% for double (1:1) occupancies based on *λ_1_* = 0.83 and *λ_2_* = 0.58 (Figure 2E). Traps with symmetric pairings, *k_1_*:*k_2_* = 1:2 and 2:1, each represented about 4% of the total, i.e., *n_t_*(1:2) = 36, *n_t_*(2:1) = 42, whereas fewer than 2% contained multiplets with *k_1_*:*k_2_* = (≥2:≥2). About 22% of the traps were empty, i.e., *k_1_*:*k_2_* = 0:0, *n_t_*(0:0) = 230. The remaining traps, classified as controls, contained cells of only one type, i.e., *k_1_*:*k_2_* = 0:≥1 (*n_t_* = 211) and ≥1:0 (*n_t_* = 146). Random trapping of fluorescently labeled U87^GFP^ cells with the NK92^IL2^ cells visually confirmed these distribution patterns (Figure 2F). A single experiment enables a wide variety of E:T ratios. The number of empty, singletons, and multiplets for particles (*λ_1_* = 0.51 and *λ_2_* = 0.58) and for cells (*λ_1_* = 0.83 and *λ_2_* = 0.58) is represented in **Table S1**.

### 2.2 Immune cell–tumor interactions

#### 2.2.1 Functional response of PBMCs against U87^GFP^ cells

PBMCs were co-cultured inside our CellTrap device with U87 glioblastoma cells (U87^GFP^) to evaluate immune–cancer cell interactions (**Figure 3A-3C and Movie S1**). The response of U87^GFP^ cells, in the presence and absence of PBMCs, was monitored by tracking their fluorescence intensity over a 14-hour period. As the PBMCs interacted with the U87^GFP^ cells, the fluorescence signal gradually weakened and eventually disappeared, indicating damage to the cancer cells (Figure 3A). However, this diminishing fluorescence effect was not observed in the control group of U87^GFP^ cells, which were not co-incubated with PBMCs (Figure 3B). Representative examples of PBMCs interacting with U87^GF^ cells at different E:T ratios highlight a heterogeneous response (Figure 3C). For example, at an E:T ratio of 1:1, cancer cell membrane integrity was compromised after 8 h, indicating that the cell is moving towards an apoptotic state, even though the green fluorescence signal has not yet completely diminished. For E:T ratio of 1:2, one cancer cell was neutralized by the immune cell after 12 h, whereas the second cancer cell migrated away from the immune cell. For an E:T ratio of 2:1, in the presence of two immune cells, the membrane of the single cancer cell was compromised after 8 h, as the cell initially transformed into a bloated phenotype before completely losing the fluorescence signal. For an E:T ratio of 2:2, both cancer cells were neutralized by 12 h, whereas a clear lag in their degree of apoptosis was apparent from the varying fluorescence signal. As an on-chip control, cancer cells maintained in the absence of PBMCs remained morphologically intact and retained stable fluorescence intensity throughout the observation period (Figure 3C).

**Figure 3.**
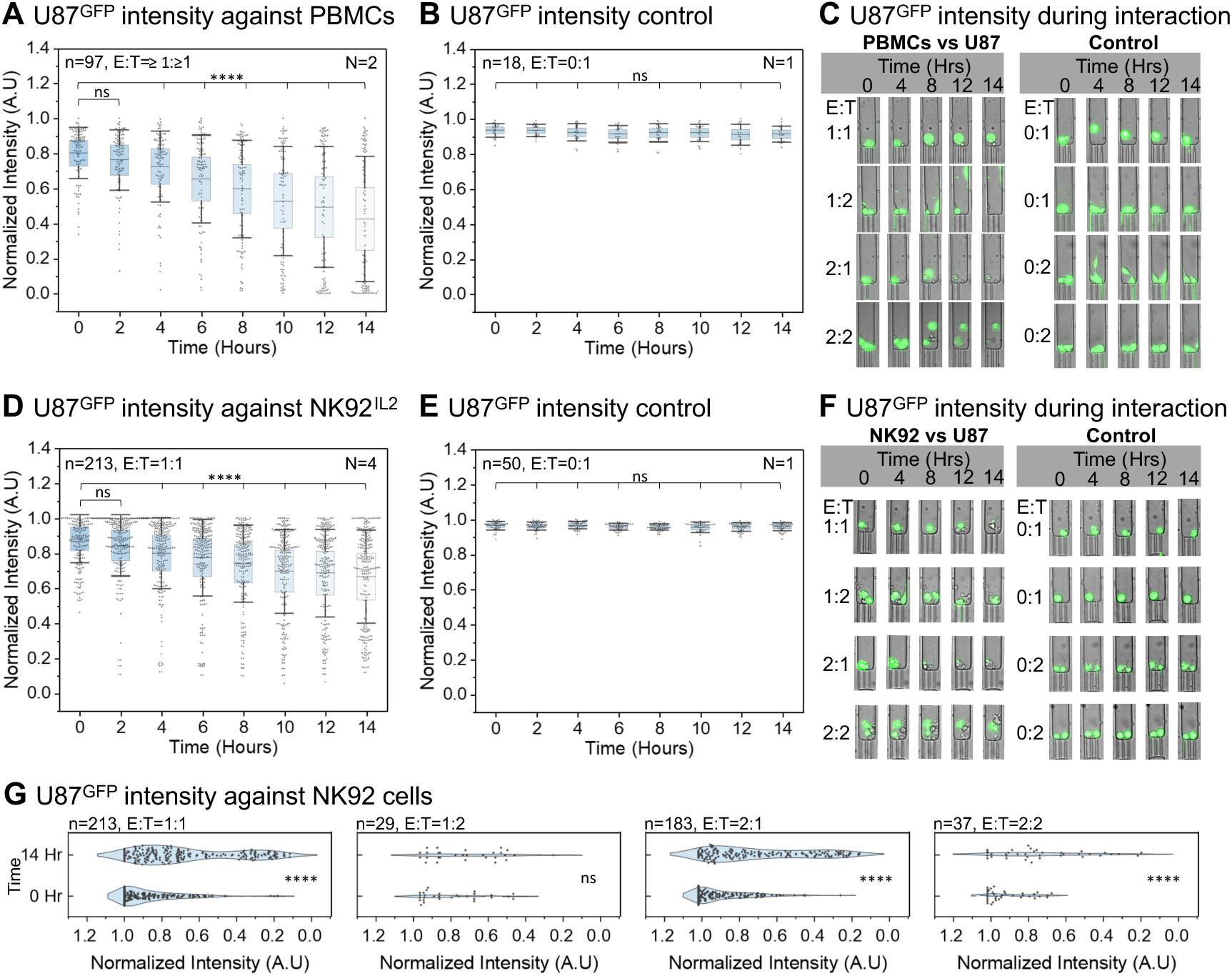
Response of PBMCs and NK92^IL2^ against U87^GFP^ cells. (A) Fluorescence intensity of U87^GFP^ cells decreases significantly after 4 h of co-incubation with PBMCs at E:T = ≥1:≥1. This data is curated from two independent CellTrap devices (N = 2), where, in total, 97 traps were analyzed (n = 97). (B) Inside one of the CellTrap devices used in (A), control traps with only one U87^GFP^ cell per trap, i.e., E:T = 0:1, were analyzed, maintaining fluorescence signal over 14 h (N = 1, n = 18). (C) Representative images of U87^GFP^ cells interacting with PBMCs at different E:T ratios, along with the control group containing only U87^GFP^ cells. (D) Fluorescence intensity of U87^GFP^ cells decreases significantly after 4 h of co-incubation with NK92^IL2^ at E:T = 1:1. This data is curated from four independent CellTrap devices (N = 4), where, in total, 213 traps with E:T = 1:1 were analyzed (n = 213). (E) Inside one of the CellTrap devices used in (D), control traps with only one U87^GFP^ cell per trap, i.e., E:T = 0:1, were analyzed, maintaining a fluorescence signal over 14 h (N = 1, n = 50). (F) Representative images of U87^GFP^ cells interacting with NK92^IL2^ at different E:T ratios, along with the control group containing only U87^GFP^ cells. (G) Fluorescence intensity of U87^GFP^ at 0 h and 14 h of co-incubation with NK92^IL2^ at E:T = 1:1, 1:2, 2:1 and 2:2. The intensity drop is significant in all E:T ratios except 1:2. This data is curated from the same CellTrap devices used in (D).

#### 2.2.2 Functional response of NK92^IL2^ cells against U87^GFP^ cells

Initially, PBMCs were used as effector cells against U87 targets to model a physiologically relevant immune response. However, due to the inherent heterogeneity and donor-dependent variability in PBMC populations, we subsequently employed the NK92^IL2^ cell line to achieve more reproducible and mechanistically interpretable results (Figure 3D-3G and **Movie S2**). This transition from PBMCs enabled precise analysis of NK92^IL2^-cell–mediated killing dynamics under controlled E:T ratios within our microfluidic CellTrap device. In this experimental campaign, we also performed viability and proliferation studies to ensure the biocompatibility of our platform under various control conditions. It was observed that during a 14-hour period, cells proliferated and maintained a viability of more than 95% (Figure S4). The interaction between NK92^IL2^ and U87^GFP^ cells at a 1:1 E:T ratio showed a statistically significant reduction in the fluorescence intensity of the cancer cells after 4 hours (Figure 3D). Moreover, a significant downward shift in the U87^GFP^ cell population, accompanied by a decrease in average fluorescence intensity from 0.88 a.u. to 0.66 a.u., was evident after 14 hours, indicating an average of nearly 22% cancer cell death. Comparison with the control group demonstrated that immune cells exert a significant killing effect on cancer cells (Figure 3E). The effect of varying E:T ratios, i.e., NK92^IL2^ cells to U87^GFP^ cancer cells, highlights a similar heterogeneous response as that observed with PBMCs vs. U87 cells (Figure 3F). At an E:T ratio of 1:1, loss of cancer cell membrane integrity was observed after 12 h of co-incubation with an immune cell. For an E:T ratio of 1:2, one of the two cancer cells was eliminated by NK92^IL2^ cells after 8 h, whereas the second cell remained viable throughout the experiment until 14 h. For an E:T ratio of 2:1, cancer cell lysis occurred earlier, at 4 h. At a 2:2 ratio, one cancer cell was neutralized after 8 h, while the other persisted until 14 h. As an on-chip control, U87^GFP^ cells maintained in the absence of NK92^IL2^ cells remained morphologically intact and retained stable fluorescence intensity throughout the 14-hour observation period. A significant population shift in U87^GFP^ cells was observed after 14 hours of incubation with NK92^IL2^ cells across different E:T ratios, except at an E:T of 1:2 (Figure 3G). In particular, E:T ratios of 1:1 and 2:1 were associated with a marked reduction in U87^GFP^ fluorescence intensity, indicative of substantial target cell killing. At E:T ratio of 1:2, a discernible shift toward reduced U87^GFP^ intensity was also observed in conditions where at least one cancer cell underwent cell death. However, the change is still not significant when accounting for both target cells within a trap, which could be due to the limited number of data points (n = 29).

#### 2.2.3 NK92^IL2^-mediated cytotoxic response against K562, LS174T, and U87 cells

##### 2.2.3.1 Short-term calcium responses

One of the earliest activation events in immune cells upon recognizing a target cancer cell is a transient increase in intracellular calcium, which is essential for cytokine secretion and cytotoxic granule-mediated killing of cancer cells^33–35^. To evaluate this calcium response, NK92^IL2^ cells are loaded with the Fluo-4 calcium imaging reagent. The labeled NK92^IL2^ cells are then incubated either alone or in co-culture with cancer cells at varying E:T ratios, and their interactions are recorded every 10 seconds for 30 minutes using an environmentally controlled fluorescence microscope (**Figure 4A and Movie S3**). When incubated with U87 cells, NK92^IL2^ cells exhibited a gradual rise and fall in calcium flux, with single-cell traces (grey lines) showing transient peaks that were largely absent in NK92^IL2^-only controls (cyan). This slower decline may reflect differences in the ligands presented by U87 cells, the stability of the immune synapse they form with NK92^IL2^ cells, or the duration of Ca²⁺ signaling engagement. In contrast, incubation with LS174T cells produced a mixture of calcium responses in NK92^IL2^ cells, whereas K562 targets induced a pronounced, sudden spike in calcium flux in immune cells. These differences in calcium peak dynamics likely reflect distinct modes of NK92^IL2^ signaling and activation upon contact with each cancer type. Because K562 cells lack MHC class I, NK92^IL2^ recognition follows the “missing-self” principle. In contrast, U87 and LS174T cells express MHC class I, so NK92^IL2^ activation depends primarily on the balance between activating and inhibitory receptor engagement, with both target types presenting higher levels of activating than inhibitory ligands^36^. Control traces (cyan) remained flat in all cases, confirming that calcium flux transients in the immune cells were specifically induced by the engagement with the target cells.

**Figure 4.**
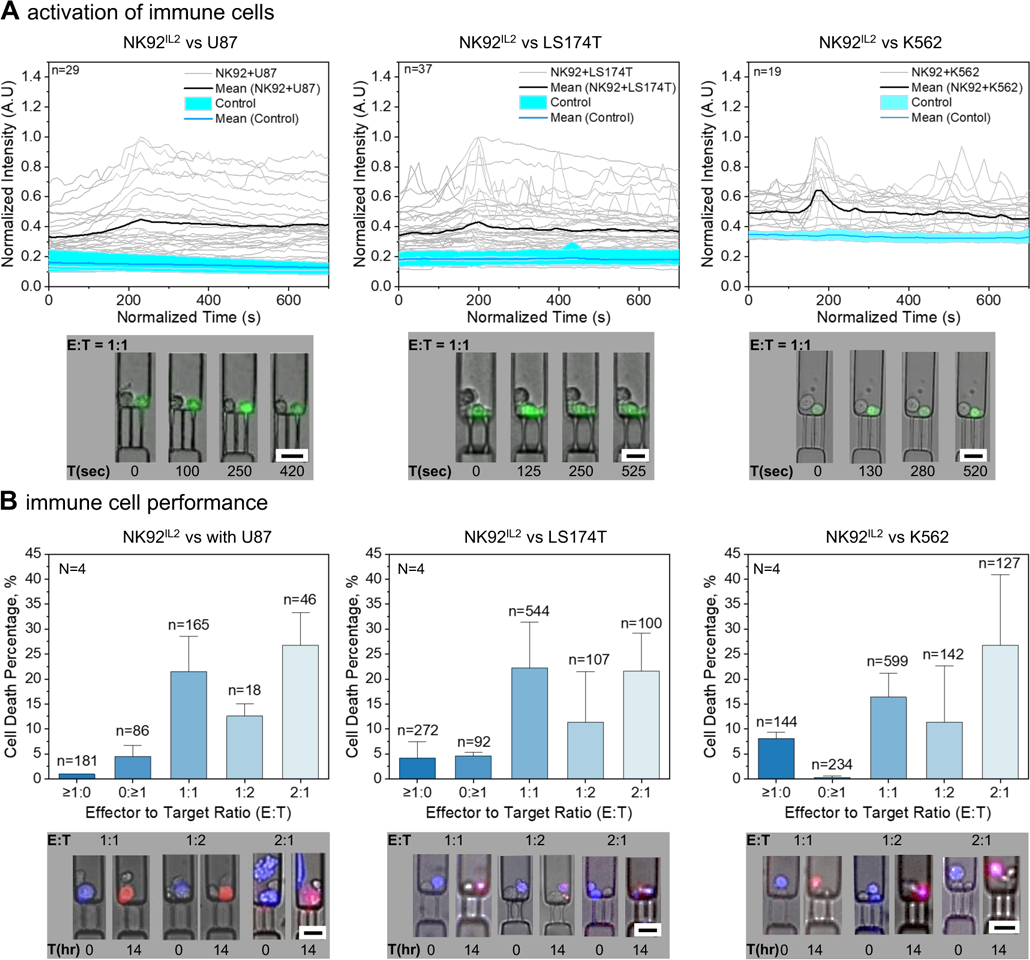
Calcium flux and killing response of immune cells against cancer cells. (A) Calcium flux (normalized intensity) in NK92^IL2^ immune cells varies over time in the presence of various cancer cell lines (U87, LS174T, K562). NK92^IL2^ cells alone show a flat response (control). Each grey line represents a single immune cell tracked. Representative images show NK92^IL2^ cells (Green) and cancer cells (U87, LS174T, K562) co-incubated in the CellTrap chip at an E:T ratio of 1:1. n = number of traps analyzed. (B) The killing response of NK92^IL2^ cells at different E:T ratios (1:1, 1:2, 2:1) against cancer cells (U87, LS174T, K562) is quantified and compared with control groups with E:T ratios of ≥1:0 and 0:≥1. N = number of CellTrap chips analyzed. n = number of traps analyzed. Representative images show NK92^IL2^ cells interacting with cancer cells (U87, LS174T, K562 in blue) with varying E:T ratios at 0 and 14 hours. Red color indicates cell death at 14 hours. Scale bars: 25µm.

##### 2.2.3.2 Extended cytotoxicity response dynamics

The immune response was further evaluated within the CellTrap device by using the NK92^IL2^ immune cells against three cancer cell lines: U87, LS174T, and K562 (Figure 4B). The effector NK92^IL2^ cells were seeded into the device, with or without target cells (K562, LS174T, or U87). After 14 hours of co-incubation, cytotoxicity was assessed using a propidium iodide (PI) assay. The killing efficiency of NK92^IL2^ cells was compared across different E:T ratios (≥1:0, 0:≥1, 1:1, 1:2, and 2:1). As expected, the percentage of target cell death increased with higher effector cell counts, indicating a dose-dependent killing response.

To rule out the cytotoxicity effect from the device, we evaluated the biocompatibility of the trapping device by focusing on cell viability and proliferation (**Figure S4**). For this, immune and cancer cells were incubated individually and in a mixed state within the trapping chip. When cells are seeded individually (either only immune cells or only cancer cells), this condition is referred to as the separate chip control. In contrast, when immune and cancer cells are mixed and seeded together in the same chip, resulting in several traps containing only one cell type, this condition is referred to as the same chip control. For both control setups, cell proliferation and viability are assessed. Trapped cells in each chamber are monitored under a microscope for 14 hours. The same-chip and separate-chip controls demonstrate similar cell viability and proliferation (Figure S4), highlighting that cell death in traps containing both immune and cancer cells (Figure 4B) results from immune-mediated killing, rather than spontaneous cell death, due to neighboring traps or any effect of PI.

For U87 cells co-incubated with NK92^IL2^ cells at an E:T ratio of 1:1, we observed an average cancer cell death of ∼21.5%, representing optimal cell–cell interaction and effective immune synapse formation that enabled this killing response. The data were averaged from four independent chips and multiple cell traps with the same E:T ratio to improve statistical confidence. When the E:T ratio was reduced to 1:2, the cytotoxicity response decreased by nearly half, to ∼12.6% cancer cell death, indicating that the limited availability of effector cells restricted NK92^IL2^ engagement and lytic granule delivery to U87 cells. Conversely, increasing the E:T ratio to 2:1 enhances the cell death rate to ∼26.7%, suggesting that higher effector density improves the probability of contact between immune-cancer cells and promotes cumulative target lysis^37^. This increasing trend in U87 cells’ death by NK92^IL2^ cells with increasing E:T ratio is consistent with bulk assays, suggesting that NK92^IL2^ cells do not immediately engage with a new cancer cell after killing the first target ^30^. We observed a cell death of ∼1% for the effector-only (≥1:0) and ∼4.4% for the target-only (0:≥1) control groups in the same chip, which confirms minimal spontaneous apoptosis and target-specific NK92^IL2^-mediated killing. Overall, the results indicate a dose-dependent killing response of NK92^IL2^ cells against U87 targets, where cytotoxicity increases with the number of effector cells until reaching near-saturation at higher E:T ratios, consistent with contact-dependent NK cell–mediated cytotoxic dynamics.

For LS174T cells co-incubated with NK92^IL2^ cells at an E:T ratio of 1:1, the cancer cell death peaked at ∼22%, which decreased to ∼11.3% for an E:T ratio of 1:2, in a response similar to NK92^IL2^-U87 cells’ interaction. However, increasing the E:T ratio to 2:1 did not significantly enhance the killing response (∼21.4%) compared to the 1:1 ratio. The cell death in the control groups, i.e., ∼4.2% in effector-only (≥1:0) and ∼4.6% in target-only (0:≥1), was significantly lower than the NK92^IL2^ cells-mediated cytotoxicity of LS174T cells. Similarly, the K562 cells co-incubated with NK92^IL2^ cells at E:T ratios of 1:1, 2:1, and 1:2 resulted in ∼16.4%, 27% and 11.3% cell death, respectively. The control groups with E:T of ≥1:0 and 0:≥1 showed cellular apoptosis of ∼8% and <1%, respectively.

The co-incubated cancer cells with NK92^IL2^ responded differently depending on the E:T ratios. For example, the NK92^IL2^ were relatively more aggressive towards the U87 (∼21.5%) and LS174T (∼22%) cells compared to the K562 (∼16.4%) when co-incubated with the cancer cells at a 1:1 ratio. At an E:T ratio of 2:1, this aggressive cytotoxicity was upregulated by ∼10.6% for K562 and ∼5.2% for U87 cells, whereas it plateaued for LS174T cells. The CellTrap platform not only enables these variable E:T ratios but also provides control group data within the same chip, which helps clarify the interpretation of cellular interactions.

## 3. Conclusions

This work presents a microfluidic cell-trapping device that enables long-term imaging and quantitative analysis of immune–cancer cell interactions at single-cell resolution. By maintaining continuous medium flow during loading, minimizing crosstalk between neighboring traps, and allowing precise control over effector-to-target (E:T) ratios, the platform addresses key limitations of conventional cytotoxicity assays that rely on bulk, endpoint measurements. The CellTrap device offers a simple, user-friendly workflow with the potential for instrument-free operation, enabling rapid cell loading within ∼5 minutes and immediate observation of cellular interactions. Stable time-lapse imaging was achieved for up to 14 hours, while maintaining viability of>90% in control groups, thereby strengthening confidence that the measured effects arise from biological processes rather than device-induced stress. Because individual cancer cells are tracked over time, cell death can be directly attributed to immune-mediated killing. The same chip simultaneously provides variable E:T ratios alongside matched on-chip controls, reducing batch effects and enabling straightforward comparisons. To support high-throughput experiments, a custom MATLAB pipeline was developed to analyze 1024 traps in parallel, allowing scalable quantification of interaction dynamics and cytotoxicity kinetics. Using NK92^IL2^ cells co-incubated with K562 and U87 cancer cells, we observed strong cytotoxicity at higher E:T ratios, while responses against LS174T cells remained comparatively unchanged; at 1:1 E:T ratios across all tested cancer types, killing efficiency was consistently ∼20–25%. Importantly, early calcium signaling and long-term fluorescence loss provided complementary readouts, capturing both rapid immune activation and delayed cytotoxic outcomes. Overall, this platform is particularly valuable in scenarios where continuous monitoring and temporal resolution are essential and where FACS-like endpoint assays are insufficient to capture heterogeneity, timing, and interaction history. Finally, the non-sealed trap design enables perfusion and multiple medium exchanges, opening the door to sequential perturbations and experimental designs that are challenging to implement with conventional droplet-based segmentation approaches. Adaptation for patient-derived cells could further establish this device as a versatile platform for personalized immunotherapy screening and mechanistic studies of tumor–immune dynamics.

## 3. Materials and methods

### 3.1 Device design

We have designed a multistage (*S*) bifurcated microchannel with a number (*N*) of cell traps defined as *N* = 4 × 2*^S^*, where the channel, after the last bifurcation, further splits into four sub-channels terminating at the cell traps. For a device with eight bifurcation stages, i.e., *S* = 8, we can calculate the total number of traps as *N* = 4 × 2^8^ = 1,024. The design is motivated by uniformly splitting a channel into two at each bifurcation stage to equal the hydraulic flow resistance (*R_h_*) in each sub-branch. The 1,024 parallel traps, with a width (*W_c_*) and gap of 30 μm each, were unfirormally spaced in a one-dimensional array spanning over 61.44 mm length of the chip, which can be conveniently fitted on a glass slide (25 mm × 75 mm). Each 30 μm wide trap has two or three filter channels with a width (*W_f_*) of 2 μm and a gap of 10 μm. The filter height (*H_f_*) of 4 μm is designed to be smaller than the channel height (*H_c_*) of 40 μm, facilitating cell trapping. The height of these two layers will be controlled during the spin-coating step of the device fabrication.

### 3.2 Device characterization

The device design is experimentally and numerically investigated for flow distribution and shear stress produced within the channels under two operational conditions, i.e., syringe pump- and pipette-driven flow. To obtain the accurate boundary conditions for the numerical simulations, we first evaluated the hydraulic flow resistance (*R*) of the designed CellTrap device experimentally as 1.48 × 10^13^ Pa.s/m^3^ (**Figure S5**). Since the flow in the CellTrap device is driven by hydrostatic pressure produced by the liquid column in a 1 mL pipette tip at the device inlet, we first evaluated the inlet pressure. For a liquid column height (*h*) of approximately 10 cm, the inlet pressure is computed as follows: *P = ρgh =* 998 Pa, where *ρ* is the density of the aqueous medium, and *g* is the gravitational acceleration. Using a known inlet pressure (*P =* 998 Pa) and measured hydraulic resistance (*R* = 1.48 × 10^13^ Pa.s/m^3^) of the CellTrap device, we calculated from the Hagen–Poiseuille relation a total flow rate through the device as *Q* = *P*/*R* = 6.74 × 10^-11^ m^3^/s = 4.04 μL/min. Therefore, it takes ∼2 min to load a concentrated cell suspension of ∼10 µL from the pipette tip to the CellTrap device. We further estimated the flow rate through the last bifurcated channel, which opens into four parallel traps, by dividing the total flow rate by 256 channels as follows: 4.04 μL/min ÷ 256 = 0.0158 μL/min. For the comparison of pump- and pipette-driven flows, the cell loading experiments in Figure S2 were performed using a total pump-driven flow rate of 5 µL/min, which translates to nearly the same value of 0.0195 µL/min of flow after the last bifurcation. This served as a reference flow rate for the inlet boundary conditions in the numerical simulations in Figure S1 and **Figure S6**.

The cell traps are simulated in COMSOL Multiphysics 6.2, using the Laminar Flow Module, for syringe pump- and pipette-driven flow conditions to evaluate flow distribution (Figure S1) and shear stress (Figure S6) when the traps are empty or filled with cells. For the pump-driven flow, the inlet boundary condition was volumetric flow rate, ranging from 0.019 to 0.215 µL/min. After each simulation, the inlet pressure was evaluated for the given flow rate boundary condition and the occupancy situation of the trap. The evaluated inlet pressure values, ranging from ∼100 to ∼6,000 Pa, served as inlet boundary conditions to simulate the respective pipette-driven flows. It was observed that the pump consistently generates a higher shear stress within the filters compared to a pipette-driven flow, particularly when the traps are occupied. This observation suggested that the pump, as the active source, continues to push the flow, regardless of the increased resistance of an occupied filter, and maintains the fixed flow rate. A syringe pump-driven flow can affect cell viability as the cells are constantly forced through the filters under high shear stress, where they can occasionally squeeze through the filters to escape the trap (**Figure S7**). On the other hand, in the pipette-driven flow, the inlet pressure stays consistent irrespective of the trap occupancy. Once a trap is occupied, the flow through this filter is significantly reduced along with the shear stress on the occupied cells; therefore, the cells are less prone to escaping the traps. The total flow rate in the device increases only when the column height in the pipette tip is increased, thereby increasing the shear stress within the channels.

### 3.3 Device fabrication

A negative photoresist (SU8-3005; Micro Resist Technology, Germany) was spin-coated on a 4-inch silicon wafer using a two-step program to ensure a uniform coating. First, the wafer was spun at 500 rpm for 10 s with an acceleration of 100 rpm/s, followed by 4500 rpm for 30 s with an acceleration of 300 rpm/s (expected layer thickness of ∼4 µm). The coated wafer was then soft-baked at 95 °C for 2 min. After soft baking, the hotplate was switched off to allow the wafer to cool gradually to room temperature, thereby minimizing thermal stress. A CAD mask of filters, including alignment markers, was prepared in AutoCAD and converted to an instrument-compatible GDS format. An ultraviolet (UV) exposure (365 nm) of the first spin-coated photoresist layer was performed using a maskless laser writer (µMLA, Heidelberg Instruments, Germany) at a dose of 180 mJ/cm². After UV exposure, the wafer was post-baked for 1 min at 65 °C and 1 min at 95 °C, then allowed to cool to room temperature on the hotplate. Next, a second negative photoresist (SU8-3025; Micro Resist Technology, Germany) was applied on top of the first layer using a two-step spin-coating process (500 rpm for 10 s at 100 rpm/s, then 3000 rpm for 30 s at 300 rpm/s with an expected thickness of the second layer as ∼25 µm). The wafer was soft-baked at 95 °C for 12 min and cooled to room temperature. The CAD mask of the main channels was aligned to the UV-exposed filters in the first layer using the alignment markers. A second UV exposure was carried out at a dose of 100 mJ/cm². The wafer was then post-baked for 1 min at 65 °C and 4 min at 95 °C, followed by gradual cooling to room temperature. Finally, the wafer was developed for 6 min in mr-Dev 600 (Micro Resist Technology, Germany). The heights of the filters and the main channels were measured as ∼4 µm and ∼35 µm, respectively, using a profilometer (Dektak XTR, Bruker Corporation, Billerica, MA, USA). For soft lithography, polydimethylsiloxane (PDMS) was prepared by mixing the elastomer and curing agent at a 10:1 ratio. The mixture was poured onto the SU-8 master mold, degassed under vacuum, and cured at 65 °C for 2 h. After curing, the PDMS layer was peeled from the mold, and inlet and outlet holes were created using a 1.2 mm biopsy punch. The PDMS device was then plasma-bonded to a glass slide and placed in an oven at 65 °C for 20 min to strengthen the bond.

### 3.4 Device operation

The device was designed to enable instrument-free and user-friendly operation. The CellTrap device operates solely via a hydrostatic pressure gradient and can therefore be driven using a standard 1 mL pipette tip. Prior to use, the chip was decontaminated with ethanol, thoroughly rinsed with aqueous media, and degassed to remove trapped air bubbles from the microchannels.

For continuous analysis of cell viability during long-term live-cell imaging, cells were seeded into the CellTrap device in RPMI 1640 culture medium supplemented with propidium iodide (PI). PI (1.0 mg/mL stock solution) was diluted 1:40,000 in RPMI 1640. Concentrated cell suspensions were prepared by resuspending each cell type in 50 µL of medium at a density of 100,000 cells/mL. The PI-containing RPMI medium was first loaded into the pipette tip, after which 20 µL of the cell suspension (≈2000 cells) was aspirated into the same tip by rotating the pipette dial anticlockwise. In this configuration, the PI-containing medium acts as a reservoir above the cell sample, preventing dilution or prolonged suspension of the cells prior to trapping. This setup minimizes the required cell volume and enables rapid and efficient seeding supported by the liquid column height in the 1 mL tip.

Once microscopy confirmed that all cells were captured in the traps, the 1 mL pipette tip was removed and replaced with shorter pipette tips, ∼2 cm long piece cut from the 100 or 200 µL tip, containing ∼20 µL of PI-supplemented RPMI. These short pipette tips were sealed with autoclave tape to reduce evaporation during imaging and were essential to accommodate the CellTrap device within the on-stage humidity and CO_2_ control chamber. The chip was then mounted on a Leica LAS X fluorescence microscope equipped with environmental control, and time-lapse imaging was performed for 20 h with adaptive focusing applied every hour.

### 3.5 Cell isolation and culture

PBMCs were isolated from freshly collected heparinized human blood using density gradient centrifugation with Ficoll-Paque (ρ = 1.077 g/mL, Cytiva). Whole blood was diluted 1:1 with phosphate-buffered saline (PBS; without Ca²⁺ and Mg²⁺) and gently layered over Ficoll in 50 mL conical tubes. Samples were centrifuged at 500 × g for 30 minutes at room temperature, with the brake disengaged, to allow for optimal gradient separation. The mononuclear cell layer at the plasma–Ficoll interface was carefully aspirated and transferred to a new tube. Cells were washed twice with PBS supplemented with 2% fetal bovine serum (FBS) by centrifugation at 300 × g for 10 minutes. Purified PBMCs were stimulated with IL-2 (100 IU/mL) and used after 24 hours.

The human cell lines NK92^IL2^, U87, K562, and LS174T were obtained from the American Type Culture Collection (ATCC, Manassas, VA, USA).

NK92^IL2^ and K562 cells were cultured in RPMI 1640 medium (Gibco™, Thermo Fisher Scientific, Waltham, MA, USA) supplemented with 10% fetal calf serum (FCS), 1% penicillin–streptomycin (Pen/Strep, Thermofisher), and 1% glutamine (Thermofisher). U87 and LS174T cells were cultured in high-glucose Dulbecco’s Modified Eagle Medium (DMEM, GlutaMAX™ Supplement; Gibco™, Thermo Fisher Scientific, Waltham, MA, USA) supplemented with 10% (v/v) heat-inactivated fetal bovine serum (FBS; Sigma-Aldrich, Merck KGaA, Darmstadt, Germany) and 1% (v/v) penicillin–streptomycin (10,000 IU/mL penicillin and 10 mg/mL streptomycin; Sigma-Aldrich). All cells were maintained at 37 °C in a humidified atmosphere containing 5% CO₂.

Adherent cell lines (LS174T and U87) were maintained in complete DMEM, while suspension cell lines (K562 and NK92^IL2^) were cultured in complete RPMI medium. NK92^IL2^ were stimulated with Human IL-2 Recombinant Protein (100 IU/mL, PeproTech®, Thermo Fisher) 24 hours before interaction analysis. Adherent cells were passaged by washing once with PBS (Gibco™), followed by detachment using trypsin (Sigma-Aldrich) and incubation for 3 to 5 minutes at 37 °C. After detachment, enzymes were neutralization with fresh complete medium. Suspension cells were collected directly from culture by transferring the cell suspension to centrifuge tubes. For both adherent and suspension cultures, cells were pelleted by centrifugation using centrifuge (Hettich® ROTINA 420R, Andreas Hettich GmbH & Co. KG, Tuttlingen, Germany) at 500×g, the supernatant was removed, and cells were resuspended in the appropriate pre-warmed complete medium for downstream experiments.

### 3.6 Calcium assay

For calcium detection, NK92^IL2^ cells were counted and approximately 200,000 cells were pelleted. Fluo-4 AM staining was performed according to the calcium imaging kit protocol (Cat. No. F10489, Thermo Fisher Scientific) by preparing a 1 mL working solution of Fluo-4 AM and adding it to the NK92^IL2^ cell pellet. Cells were incubated at 37 °C for 30 min, followed by an additional 30 min at room temperature. After staining, cells were washed three times with PBS (without Ca²⁺ and Mg²⁺) by centrifugation at 500×g. The stained NK92^IL2^ cells were then split into two equal fractions: one was mixed with cancer cells (U87, LS174T, or K562), while the other served as a control with NK92^IL2^ cells only. Samples were loaded into the microfluidic chip following the device operation protocol described above. Time-lapse imaging was carried out on an environmentally controlled microscope (Leica LAS X Thunder) using a 10× objective, acquiring brightfield and fluorescence images every ∼10 s for 30 min, with control and experimental chips imaged in parallel. Image sequences were analyzed in ImageJ (https://imagej.net/ij/), and calcium flux was quantified from changes in Fluo-4 fluorescence intensity.

### 3.7 Post-processing

For the analysis of cell death, proliferation, and viability, stitched images of the 1024 traps at 0 and 14 hours were manually analyzed. The number of cancer or immune cells in a live or dead state was counted at a given time step and processed in Excel. For experiments involving fluorescently labeled U87^GFP^ cells (Figure 3), stitched images of the 1024 traps, obtained every two hours, were post-processed in MATLAB. Brightfield and fluorescence channels from the stitched images were straightened and then cropped. The stitched images were split into 1024 images corresponding to each trap within the CellTrap device. The number of cancer cells in each split image or trap was counted, and the change in fluorescence intensity over time was simultaneously analyzed. For each trap, a representative split image was saved, and intensity traces were retained only for traps containing both immune and cancer cells for subsequent analysis. The number of immune cells in the corresponding traps was manually tallied from the brightfield images to confirm the E:T ratio.

### 3.8 Statistical analysis

All statistical analyses in Figure 3 were performed using GraphPad Prism version 10 (GraphPad Software, San Diego, CA, USA). Data are presented as mean ± standard deviation (SD) from one to four independent experiments. Differences between groups were assessed using one-way analysis of variance (ANOVA) followed by Tukey (normal distribution) or Kruskal–Wallis and Dunn’s multiple comparisons test (nonparametric test). A p-value of < 0.05 was considered statistically significant. In the figures, significance levels are indicated as shown ∗p < 0.0332, ∗∗p < 0.0021, ∗∗∗p < 0.0002, and ∗∗∗∗p < 0.0001.

## Acknowledgments

This work is supported by the German Research Foundation (DFG) project #539433772, the German Academic Exchange Service (DAAD) doctoral research grant #57588370, and the TranslaTUM Seed Fund project 24-4DesWan.

---

**Figure S1.**
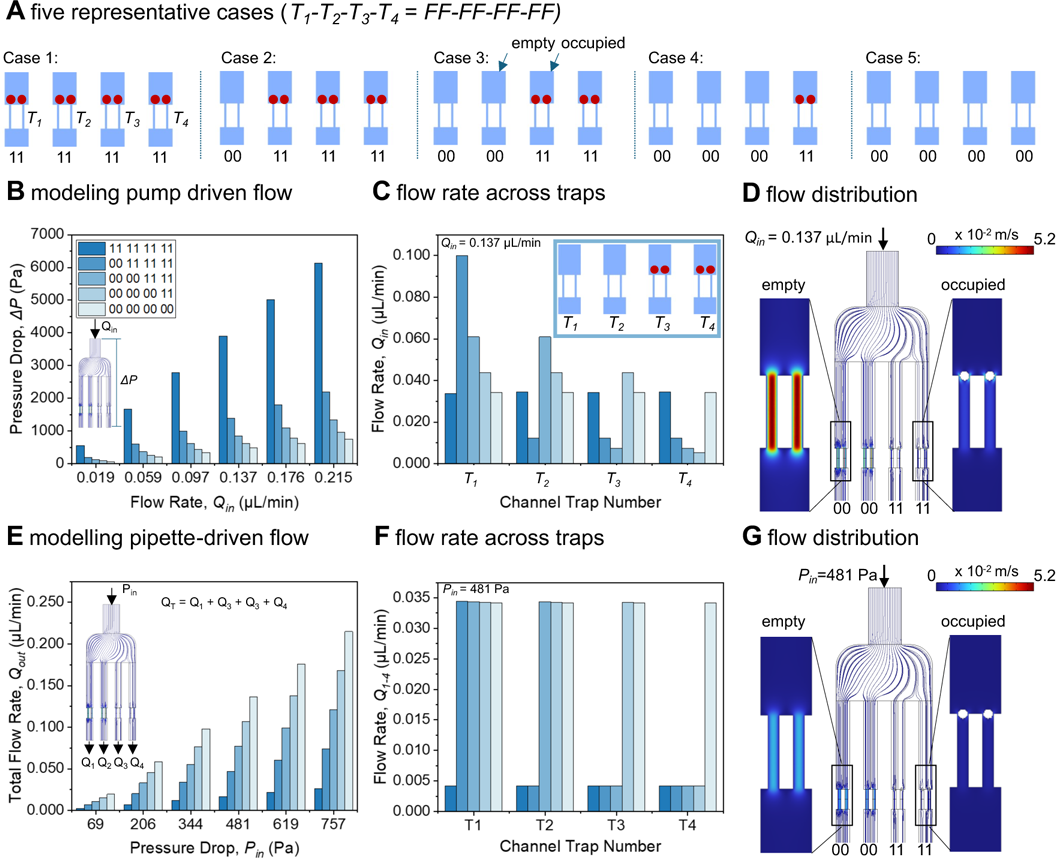
Numerical analysis of flow rate distribution within four parallel daughter channels of a CellTrap device under syringe pump and pipette-driven flow. (A) Five representative cases of traps are defined for this numerical investigation: Case 1: 11-11-11-11, Case 2: 00-11-11-11, Case 3: 00-00-11-11. Case 4: 00-00-00-11, and Case 5: 00-00-00-00. For any given case, each pair of numbers indicates the status of one of the four branches ending in an isolated cell trap (*T*), such that the case can be represented as *T_1_-T_2_-T_3_-T_4_*. Each trap (*T_1_-T_4_*) has two filters (*FF*) that can be in an open (not occupied) or closed (occupied) state; therefore, we can also define the status of the four parallel traps *T_1_-T_2_-T_3_-T_4_* as *FF-FF-FF-FF*, where *F* is equal to 0 (open) or 1 (closed). For example, in Case 1: *T_1_-T_2_-T_3_-T_4_* = 11-11-11-11, all the filters across four traps are occupied by a particle, whereas in Case 5: *T_1_-T_2_-T_3_-T_4_* = 00-00-00-00, all the filters across four traps are open. (B) For numerical modeling of the flow through the four daughter traps, we first assumed a total flow rate through the device ranging from 5 µl/min to 55 µl/min and then divided the total flow rate by 256 channels after eight bifurcations. This resulted in our inlet boundary condition, in the form of a constant flow rate ranging from 0.019 µl/min to 0.215 µl/min, which mimics an experimental syringe-pump-driven flow, for our simulations. The pressure drop (*ΔP*) across the filters increased linearly with the inlet flow rate (*Q_in_*), as predicted by the Hagen-Poiseuille equation. The highest slope for Case 1: 11-11-11-11 is associated with the highest fluidic resistance of the fully occupied traps. (C) The flow distribution through each daughter trap (*T_1_-T_4_*) was further analyzed as a function of trap occupancy by particles. A representative case was evaluated with an inlet boundary condition of *Q_in_* = 0.137 μL/min. For Case 1 (*T_1_*-*T_2_*-*T_3_*-*T_4_* = 11-11-11-11) and Case 5 (*T_1_*-*T_2_*-*T_3_*-*T_4_* = 00-00-00-00), the flow rates through all four traps remain identical because the hydrodynamic resistance is balanced across the branches. In contrast, for Case 3 (*T_1_*-*T_2_*-*T_3_*-*T_4_* = 00-00-11-11), the flow rate through traps *T_1_* and *T_2_* is higher due to lower flow resistance and the absence of particles. Consequently, branches *T_3_* and *T_4_* exhibit substantially reduced flow due to higher flow resistance and the presence of trapped particles. This indicates that, while the syringe pump supplies a constant total flow rate of 0.137 μL/min, the flow is redistributed among the individual traps according to their occupancy, such that the sum of the flow rates through all four traps remains conserved. (D) A streamline and contour representation for Case 3 (*T_1_*-*T_2_*-*T_3_*-*T_4_* = 00-00-11-11) depicts high velocity (*v_avg_* = 29.07 mm/s) for *T_1_* and *T_2_* channels and low velocity (*v_avg_* = 3.56 mm/s) for *T_3_* and *T_4_* channels. A shift in streamlines towards *T_1_* and *T_2_* channels can be clearly seen right after the flow enters from the main channel. This highlights that once a trap is occupied, it results in a diversion of flow towards open or non-occupied traps. (E) For the numerical modeling of the pipette-operated flow through the four daughter traps, we first computed the pressure drop *ΔP* across the channel for a range of total inlet flow rates *Q_in_* from 0.019 μL/min to 0.215 μL/min. This pressure drop was then used as the inlet pressure boundary condition *P_in_*, ranging from 69 Pa to 894 Pa. For each *P_in_*, the corresponding total outlet flow rate *Q_out_* (= *Q_1_* + *Q_2_* + *Q_3_* + *Q_4_*) was calculated, and in all cases, the overall flow rate increased monotonically with *ΔP*. (F) The flow distribution through each daughter trap was further analyzed as a function of trap occupancy by particles. As a representative case, we evaluated an inlet pressure boundary condition of *P_in_* = 481 Pa, which corresponds to the pressure drop *ΔP* obtained for *Q_in_* = 0.137 μL/min. The total outlet flow rate *Q_out_* (= *Q_1_* + *Q_2_* + *Q_3_* + *Q_4_*) differs significantly between two extreme cases, i.e., 0.0168 μL/min for Case 1 (*T_1_-T_2_-T_3_-T_4_* = 11-11-11-11) and 0.137 μL/min for Case 5 (*T_1_-T_2_-T_3_-T_4_* = 00-00-00-00), because there is no pump enforcing a fixed flow rate. As expected, for Case 3 *(T_1_-T_2_-T_3_-T_4_* = 00-00-11-11), the flow rate through traps *T_1_* and *T_2_* is higher due to the lower hydraulic resistance in the absence of particles, whereas branches *T_3_* and *T_4_* exhibit substantially reduced flow owing to the increased resistance from trapped particles. Unlike in the pump-driven model, each trap in the pipette-operated configuration behaves independently, and the flow through each channel is bounded between a minimum (0.0042 μL/min) and a maximum (0.0342 μL/min) value: the minimum flow rate corresponds to both filters in a trap being occupied, and the maximum flow rate corresponds to both filters being empty. (G) A streamline and velocity contour representation for Case 3 (*T_1_-T_2_-T_3_-T_4_* = 00-00-11-11) illustrates high velocities (*v_avg_* = 16.4 mm/s) in channels *T_1_* and *T_2_* and low velocities (*v_avg_* = 2.08 mm/s) in *T_3_* and *T_4_*. Similar to panel (D), a clear diversion of streamlines toward *T_1_* and *T_2_* is observed immediately after the flow enters from the main channel, indicating that once a trap is occupied, the flow is redirected toward open (unoccupied) traps. However, the *v_avg_* = 16.4 mm/s in the pipette-driven flow (G) is significantly smaller than *v_avg_* = 29.07 mm/s in the pump-driven flow (D), highlighting a low shear stress flow suitable for cell trapping without significant cell leakage through the traps. Finally, panels (D) and (G) compare device performance under the same pressure-drop conditions, where (D) corresponds to pump operation and (G) to pipette operation. For the same imposed pressure drop, the velocities in channels *T_1_* and *T_2_* of the pump-operated device are significantly higher than those in the pipette-operated device.

**Figure S2.**
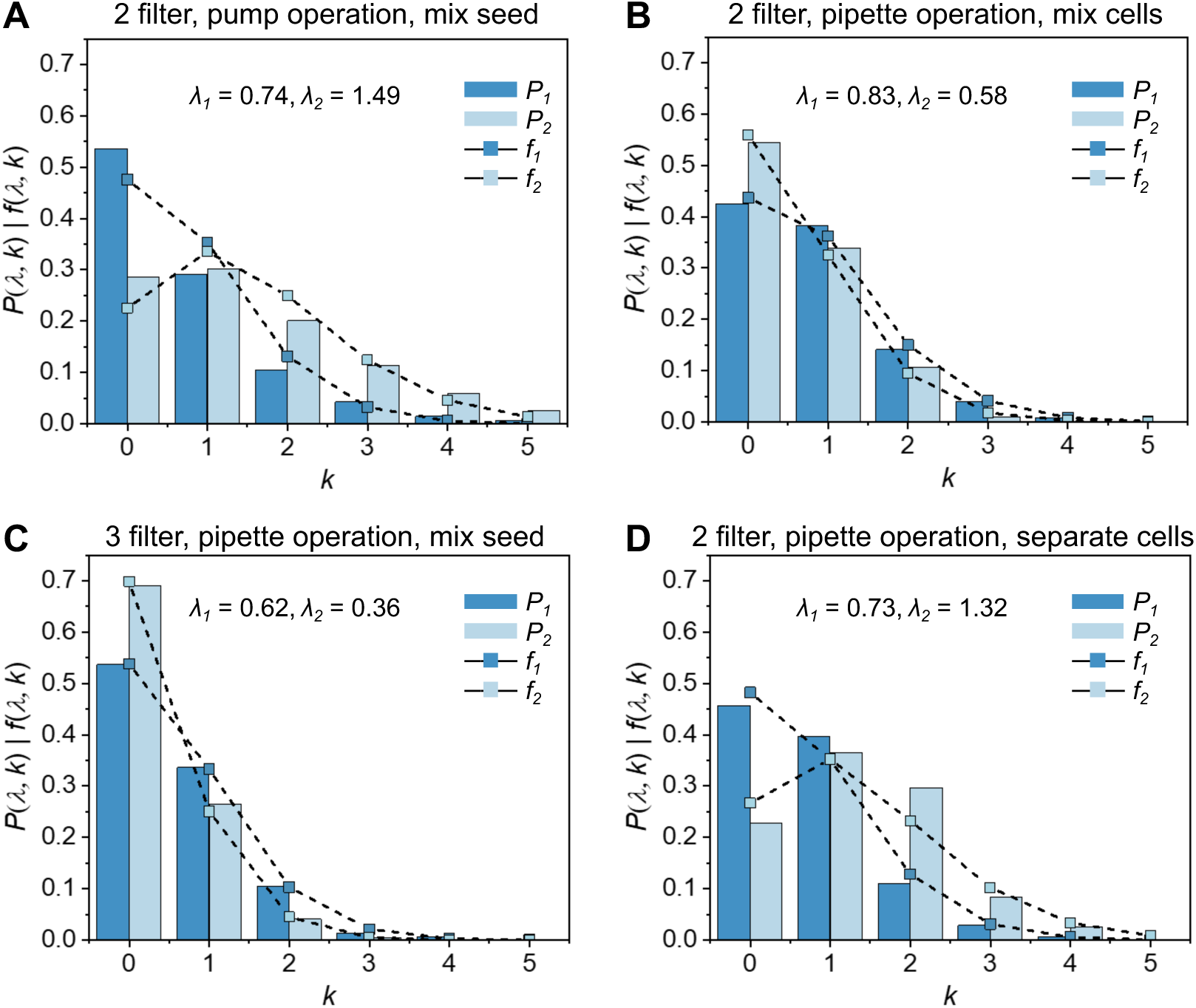
Comparison of experimental and theoretical distribution of immune (NK92^IL2^) and cancer (U87^GFP^) cells with the CellTrap device for different operating conditions. (A) A premixed sample with *λ_1_* = 0.74 and *λ_2_* = 1.49, seeded using a pump, and observed in a trapping device with 2 filter geometries. (B) A premixed sample with *λ_1_* = 0.83 and *λ_2_* = 0.58, seeded using a pipette, and observed in a trapping device with 2 filter geometries. (C) A premixed sample with *λ_1_* = 0.62 and *λ_2_* = 0.36, seeded using a pipette, and observed in a trapping device with 3 filter geometries. (D) Cancer and immune cells were seeded separately with *λ_1_* = 0.73 and *λ_2_* = 1.32, respectively, using a pipette, and observed in a trapping device with 2 filter geometries.

**Figure S3.**
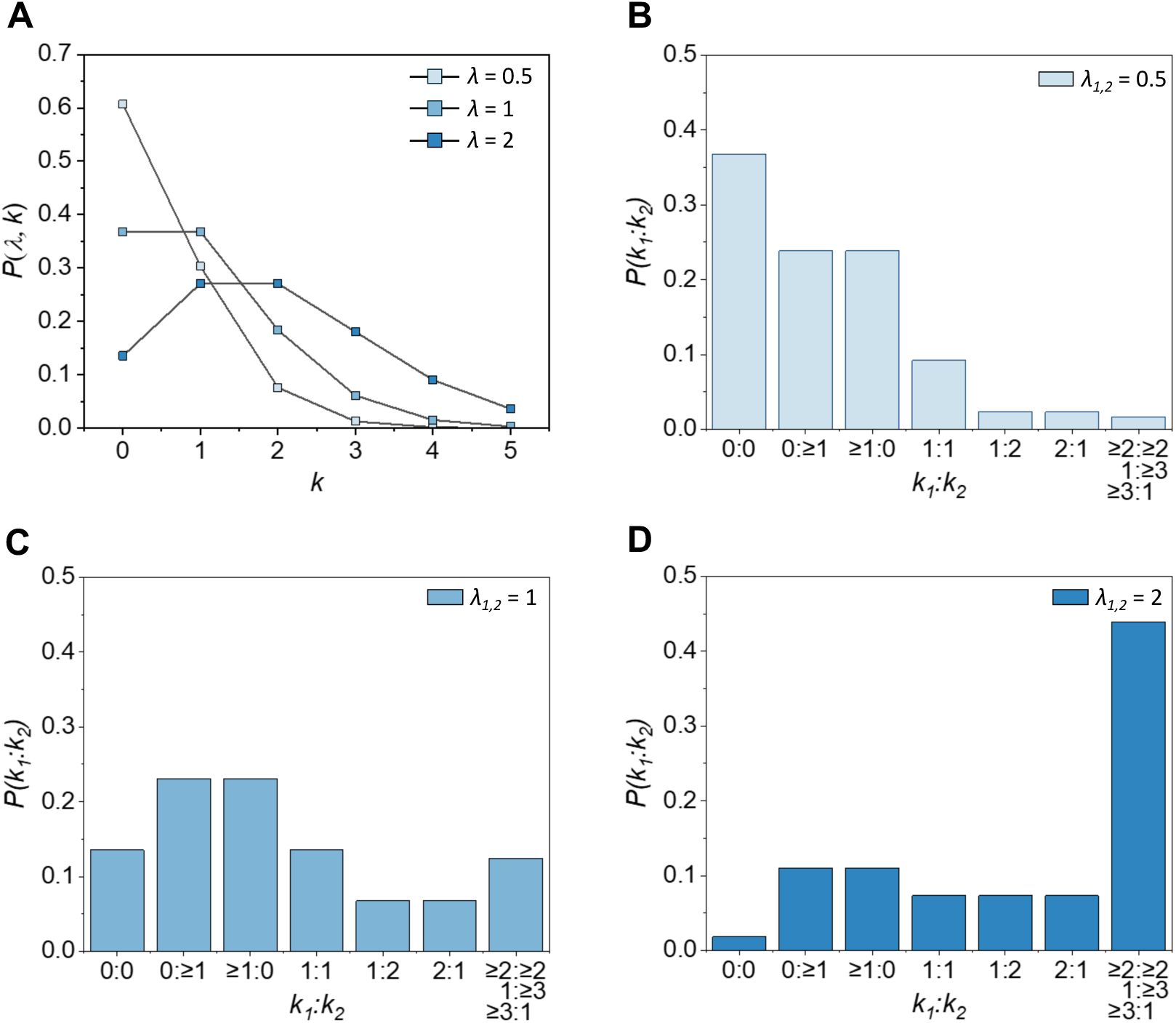
Theoretical Poisson distribution of trapped cells within the CellTrap device at different *λ* values. (A) As expected, the proportion of multiplets per trap increases with *λ* as the fraction of empty traps gradually decreases. (B-D) Double loading distribution for *λ* = 0.5, 1, and 2.

**Figure S4.**
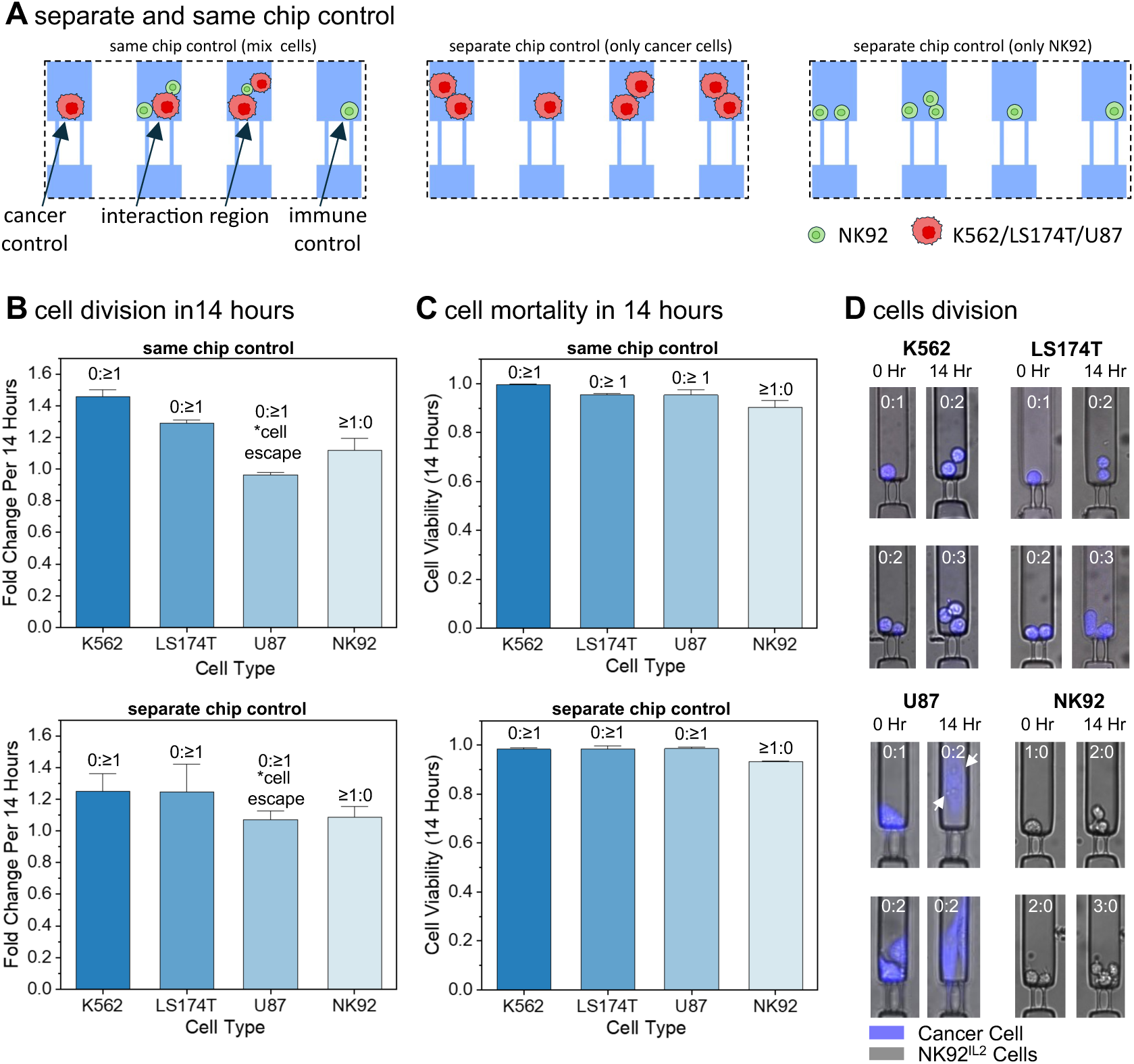
Cell proliferation and viability in separate- and same-chip controls. (A) Representative schematic of same chip control and separate chip control for cell proliferation and viability evaluation. Same chip control represents different traps containing only cancer or immune cells, while neighbouring traps in the same chip contain a mixture of immune and cancer cells. Separate chip control represents cancer cells and immune cells seeded in two separate chips. (B) A comparison of same-chip and separate-chip controls by plotting the increase in cells numbers or fold-change after 14 hours of incubation. The lowest fold-change recorded for the U87 cells can be ascribed to their tendency to escape after division, which limits cell tracking within the field of view. (C) A comparison of same-chip and separate-chip controls by plotting the cell viability (excluding divided cells) after 14 hours. The viability was >93% for all the cell types in both the control groups. (D) Pictorial representation of the cell division over the course of 14 hours of incubation obtained from the same-chip control.

**Figure S5.**
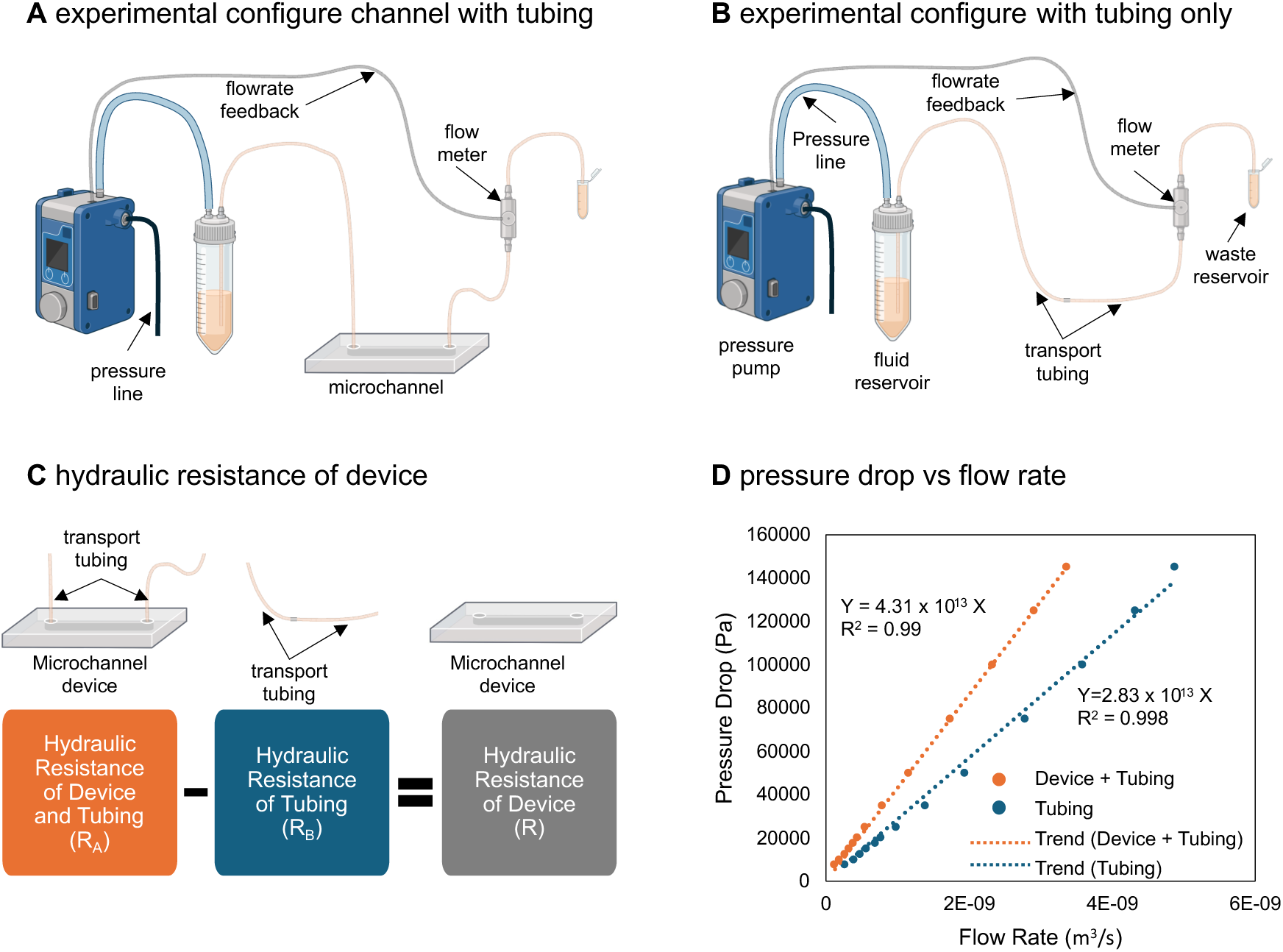
Evaluation of the hydraulic flow resistance of the CellTrap device. (A) An experimental configuration to estimate the hydraulic resistance of the device and connecting tubing for a given inlet pressure (*P*) by the pressure pump and measured outlet flow rate (*Q*) using the flow sensor. (B) An experimental configuration to estimate the hydraulic resistance of only the tubing, when the device is disconnected from the fluidic circuit, for a given inlet pressure (*P*) by the pressure pump, and measured outlet flow rate (*Q*) using the flow sensor. (C) The hydraulic resistance of the device is computed by subtracting the hydraulic resistance of the tubing from the combined hydraulic resistance of the device and tubing. (D) The inlet pressure (*P*) and the flow rate (*Q*) are plotted for configurations (A) and (B). The slopes of the trendlines are equivalent to the hydraulic resistances of the two configurations, i.e., *R_A_* = 4.31 x 10^13^ Pa.s/m^3^ and *R****_B_*** = 2.83 x 10^13^ Pa.s/m^3^. The hydraulic resistance of the CellTrap device *R* = *R_A_* - *R_B_* = 1.48 x 10^13^ Pa.s/m^3^.

**Figure S6.**
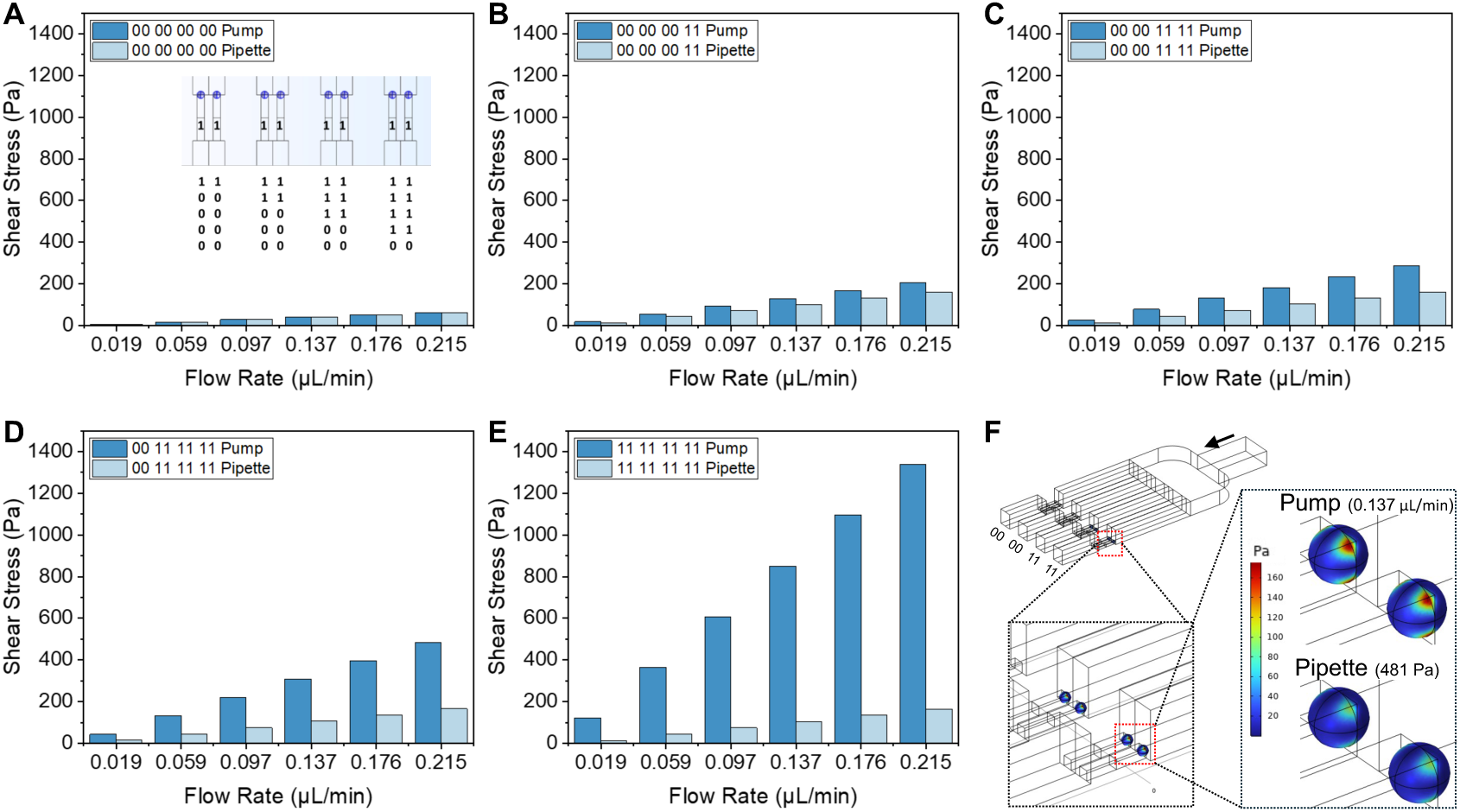
Numerical modelling of shear stress on trapped cells. (A-E) The COMSOL Multiphysics model is simulated to evaluate the maximum shear stress on trapped particles at various flow rates for both pump and pipette operations. Traps are considered under conditions of no obstruction and different particle occupancy. (F) COMSOL simulation representing shear stress at the same flow rate and pressure drop on particles obstructing the filter of traps.

**Figure S7.**
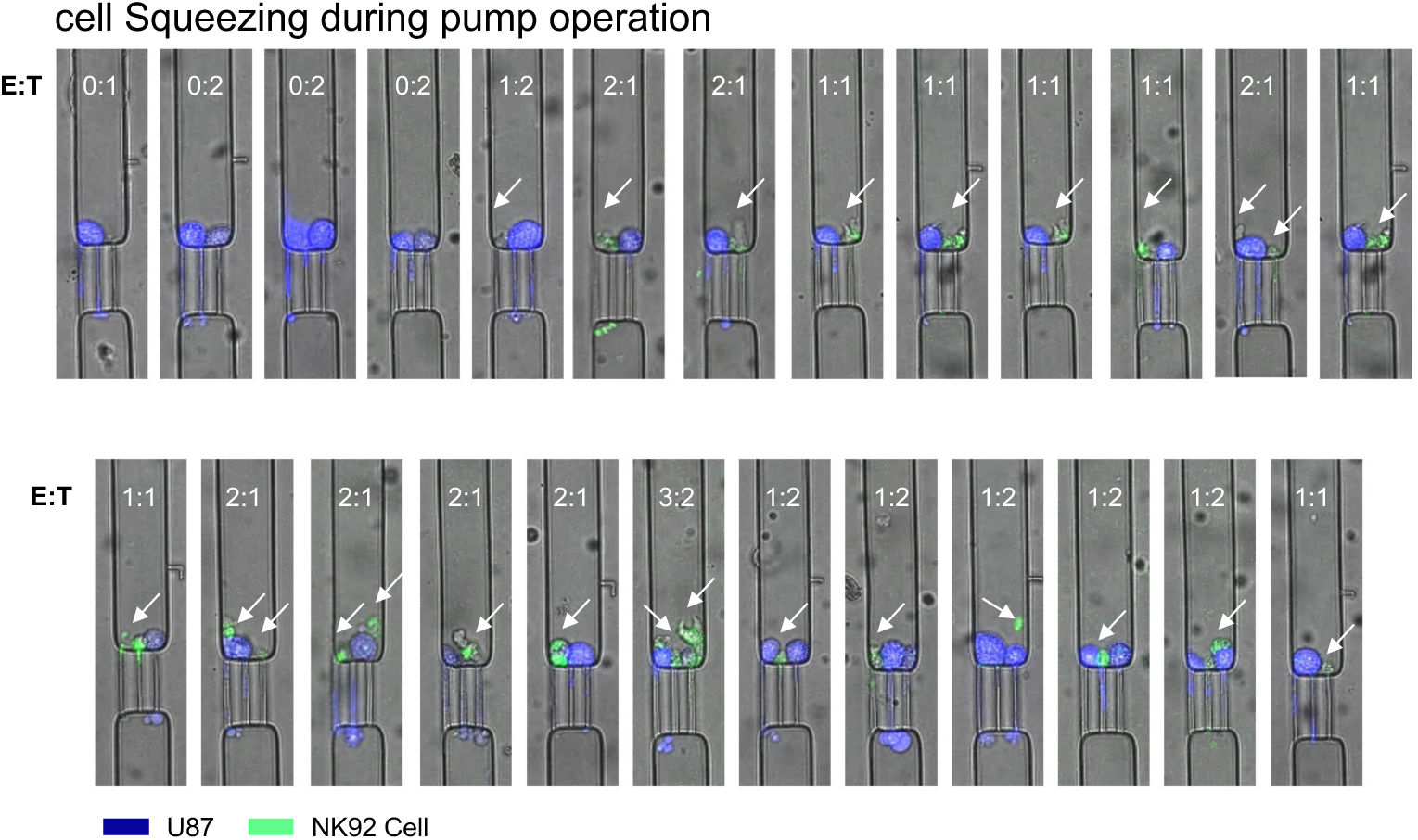
High shear effect on cells loaded using a syringe pump. U87 and NK92^IL2^ cells are stuck in the filters when the sample is loaded using the pump. Therefore, the operation using a pipette is employed in the biological sample studies. White arrow representing NK92^IL2^ cells.

**Table S1:** Detailed number of combinations of empty, cancer control, immune control, singletons at different E:T, and multiples at different E:T for particles at *λ_1_* = 0.51, *λ_2_* = 0.54 and cells at *λ_1_* = 0.83, *λ_2_* = 0.58. Data associated with Figure 2.

|  | **Effector to Target**<br>**(E:T)** | **Particles**<br>**($\lambda_I = 0.51, \lambda_2 = 0.54$)** | **Cells**<br>**($\lambda_I = 0.83, \lambda_2 = 0.58$)** |
| --- | --- | --- | --- |
| **Empty** | 0:0 | 391 | 230 |
| **Cancer control** | 0:1 | 183 | 211 |
|  | 0:2 | 49 | 92 |
|  | 0:3 | 9 | 22 |
|  | 0:4 | 0 | 4 |
| **Immune control** | 1:0 | 141 | 146 |
|  | 2:0 | 52 | 57 |
|  | 3:0 | 9 | 5 |
|  | 4:0 | 0 | 0 |
| **Singletons** | 1:1 | 105 | 142 |
| **Multiple** | 2:2 | 8 | 11 |
|  | 3:3 | 1 | 1 |
|  | 4:4 | 0 | 0 |
|  | 1:2 | 32 | 36 |
|  | 1:3 | 5 | 3 |
|  | 1:4 | 0 | 0 |
|  | 2:1 | 26 | 42 |
|  | 2:3 | 1 | 0 |
|  | 2:4 | 0 | 0 |
|  | 3:1 | 7 | 12 |
|  | 3:2 | 4 | 5 |
|  | 3:4 | 0 | 0 |
|  | 4:1 | 1 | 4 |
|  | 4:2 | 0 | 0 |
|  | 4:3 | 0 | 1 |

## Supporting information

**Movie S1: Interaction of PBMCs with U87 cells at different effector-to-target (E:T) ratios.** At an E:T ratio of 1:1, PBMCs actively engage with U87 cells, and after 5 hours, the membrane integrity of the U87 cells is lost in all five representative traps, indicating apoptosis. At an E:T ratio of 2:1, U87 cells show similar apoptotic behavior, and in addition, some PBMCs appear exhausted and dying in certain traps. At an E:T ratio of 1:2, we still observe the death of at least one U87 cancer cell. Finally, at an E:T ratio of 2:2, PBMCs can be seen killing both U87 cells. In the control group (E:T = 0:≥1), U87 cells remain viable throughout the entire imaging period.

**Movie S2: Interaction of NK92^IL2^ cells with U87 cells at different effector-to-target (E:T) ratios.** At an E:T ratio of 1:1, NK92^IL2^ and U87 cells interact within the channel, and the U87 cell can be seen dying within 12 hours. Over time, the NK92^IL2^ cells also appear to become exhausted and die. At an E:T ratio of 2:1, the increased number of NK92^IL2^ cells leads to U87 cell death within 8 hours. At an E:T ratio of 1:2, at least one U87 cell is killed by NK92^IL2^ cells by the end of the 12-hour period. In the control group (E:T = 0:≥1), U87 cells remain viable throughout the entire observation period.

**Movie S3: Calcium response of NK92^IL2^ cells interacting with U87, LS174T, and K562 cells.** The NK92^IL2^ cells against U87 cells exhibit a gradual increase in calcium signal followed by a slow decline over a period of 980 seconds. The NK92^IL2^ cells against LS174T cells show an initial rapid calcium peak followed by a decline over 980 seconds. Multiple subsequent peaks are observed, indicative of target recognition and repeated engagement. NK92^IL2^ cells against K562 cells display a rapid initial calcium peak followed by a decline over 980 seconds, similar to their response against LS174T cells. Recurring peaks are observed, reflecting ongoing target identification and attack.

